# Measurement error and resolution in quantitative stable isotope probing: implications for experimental design

**DOI:** 10.1101/2020.02.25.965764

**Authors:** Ella T. Sieradzki, Benjamin J. Koch, Alex Greenlon, Rohan Sachdeva, Rex R. Malmstrom, Rebecca L. Mau, Steven J. Blazewicz, Mary K. Firestone, Kirsten Hofmockel, Egbert Schwartz, Bruce A. Hungate, Jennifer Pett-Ridge

## Abstract

Quantitative stable isotope probing (qSIP) estimates the degree of incorporation of an isotope tracer into nucleic acids of metabolically active organisms and can be applied to microorganisms growing in complex communities, such as the microbiomes of soil or water. As such, qSIP has the potential to link microbial biodiversity and biogeochemistry. As with any technique involving quantitative estimation, qSIP involves measurement error; a more complete understanding of error, precision and statistical power will aid in the design of qSIP experiments and interpretation of qSIP data. We used several existing qSIP datasets of microbial communities found in soil and water to evaluate how variance in the estimate of isotope incorporation depends on organism abundance and on the resolution of the density fractionation scheme. We also assessed statistical power for replicated qSIP studies, and sensitivity and specificity for unreplicated designs. We found that variance declines as taxon abundance increases. Increasing the number of density fractions reduces variance, although the benefit of added fractions declines as the number of fractions increases. Specifically, nine fractions appear to be a reasonable tradeoff between cost and precision for most qSIP applications. Increasing replication improves power and reduces the minimum detectable threshold for inferring isotope uptake to 5 atom%. Finally, we provide evidence for the importance of internal standards to calibrate the %GC to mean weighted density regression per sample. These results should benefit those designing future SIP experiments, and provide a reference for metagenomic SIP applications where financial and computational limitations constrain experimental scope.

**Importance:** One of the biggest challenges in microbial ecology is correlating the identity of microorganisms with the roles they fulfill in natural environmental systems. Studies of microbes in pure culture reveal much about genomic content and potential functions, but may not reflect an organism’s activity within its natural community. Culture-independent studies supply a community-wide view of composition and function in the context of community interactions, but fail to link the two. Quantitative stable isotope probing (qSIP) is a method that can link the identity and function of specific microbes within a naturally occurring community. Here we explore how the resolution of density-gradient fractionation affects the error and precision of qSIP results, how they may be improved via additional replication, and cost-benefit balanced scenarios for SIP experimental design.

## Introduction

Stable Isotope Probing (SIP) of nucleic acids is one of the few non-culture dependent methods that can identify the functionality of microorganisms in their native environments, making it one of the most powerful techniques in microbial ecology (Radajewski et al. 2000; Manefield et al. 2002; Radajewski, McDonald, and Murrell 2003; Neufeld et al. 2007; Chen and Murrell 2010). In SIP, a substrate labeled with a heavy isotope is added to an environmental sample. Following an incubation period ranging from hours to weeks (depending on the substrate uptake rate) the DNA (or RNA) of growing microorganisms that have consumed the isotope-enriched substrate becomes more dense due to their incorporation of the heavy isotope. Community nucleic acids can then be extracted and separated in a density gradient using ultracentrifugation. DNA/RNA from organisms that incorporated the labeled substrate will appear in denser fractions of the gradient compared to where they would be with addition of an unlabeled substrate (Lueders, Manefield, and Friedrich 2004; Neufeld et al. 2007). While there has been some consideration of best practices for handling SIP data (Neufeld et al. 2007; Lueders et al., 2010; Hungate et al. 2015; Youngblut, Barnett, and Buckley 2018; Barnett, Youngblut, and Buckley 2019; Dumont and García 2019; Barnett and Buckley 2020), there remain outstanding methodological issues and questions regarding reproducibility, sensitivity and the minimum detectable effect size, some of which we address here.

A major advantage of SIP is that it can be performed on intact environmental communities, thereby taking into account microbial interactions which are missed in cultivation-based studies. Most current SIP methods require amplifying marker genes, usually 16S rRNA, from each fraction to identify substrate assimilators. However, to look at multitrophic interactions beyond co-occurrence, it is more ideal to use shotgun sequencing of whole community DNA, but the combination of SIP with metagenomic analysis quickly becomes limiting both financially and computationally. Therefore, some investigators have tried to limit shotgun sequencing either by sequencing only highly labeled fractions (Barnett and Buckley 2020), by pooling density fractions or by sequencing the unfractionated DNA and matching assembled genomes to SIP-identified substrate assimilators (Dumont et al. 2006; Murrell and Whiteley 2010; Dombrowski et al. 2016; Thomas, Corre, and Cébron 2019; Sieradzki, Morando, and Fuhrman 2019). Since metagenomic sequencing leads to financial and computational costs that are much higher than those of 16S analysis, the knowledge of the minimum number of fractions that can lead to a comparable result will be crucial.

Quantitative SIP (qSIP) is a recently developed adaptation of SIP that makes substrate uptake measurements possible at the individual or population genome scale (Hungate et al. 2015; Koch et al. 2018). In qSIP, isopycnic separation of nucleic acids in cesium chloride is combined with a mathematical model to quantify isotope enrichment. This approach allows a user to measure growth and mortality rates of individual taxa in complex communities, particularly when using ^18^O-labeled ‘heavy water’ as a substrate –since cells incorporate oxygen from water during nucleic acid synthesis, quantitatively reflecting cell division (DNA synthesis) and metabolism (RNA synthesis) (Schwartz 2007; Blazewicz and Schwartz 2011). Similarly, cell mortality rates may be quantitatively related to the degradation of unlabeled nucleic acids. By normalizing relative abundance to the total number of organisms per fraction estimated by qPCR of 16S-rRNA, qSIP has been shown to be less susceptible to taxon abundance and level of enrichment compared to other SIP methods (Youngblut, Barnett, and Buckley 2018). Hence, qSIP may be a preferred approach for combining SIP and metagenomics.

Designing qSIP experiments involves a tension between collecting many density fractions per sample (small fraction size) versus the costs of labor and sequencing. While early SIP studies inspected only the ‘heaviest’ fractions—considered to host the most isotopically enriched DNA— these fractions may contain unlabeled high GC-content DNA. The current practice is to examine many density fractions and perform statistical analyses comparing isotope-labeled versus unlabeled controls, to indicate the extent to which organisms have “shifted” within a density gradient in response to the isotope treatment (Hungate et al. 2015; Youngblut, Barnett, and Buckley 2018). Density shifts can be used to calculate substrate assimilation rate per taxon (atom % excess), and when using the universal substrate H_2_^18^O, they can be used to infer specific growth rates (Blazewicz and Schwartz 2011; Blazewicz, Schwartz, and Firestone 2014; Papp et al. 2018; Koch et al. 2018). However, even the most basic experiment (e.g one type of substrate, 2 timepoints, 3 replicates, 10 density fractions per sample) can easily generate over 100 samples for processing and sequencing. Thus, it is critical to know how to balance experimental design to ensure high quality data at sustainable costs. Doing so becomes even more important as we transition to more ambitious applications, such as metagenomics qSIP (MG-qSIP), since shotgun sequencing adds even higher costs and the amount of data per sample quickly becomes a computational limitation.

Within a replicated qSIP experiment, it is possible to evaluate statistical power - the probability of detecting a given level of isotopic enrichment. Yet, power is rarely evaluated, because, in practice, avoiding Type I errors is prioritized above avoiding Type II errors. Traditionally, many view that incorrectly inferring that a treatment is effective is more hazardous than concluding it is not effective, when in fact it is. Power analysis involves evaluating the tradeoffs among several parameters: 1) The effect size of interest, which in the case of qSIP experiments is the density shift (or amount of isotope incorporation) that the researcher wishes to detect (this can be thought of as the minimum detectable difference); 2) the acceptable α value, or acceptable probability of Type I error (for qSIP, a type I error occurs when the researcher concludes that there is isotope incorporation when in fact none occurred); 3) the acceptable β value, or acceptable probability of Type II error (for qSIP, a Type II error occurs when the researcher infers “no isotope incorporation”, when in fact some isotope incorporation actually occurred); and 4) the number of true, independent, replicates (sample size) used in the experiment. Power is defined as 1 - β. It is the probability that a true difference will be detected in a given experimental design. Applied to qSIP, power analysis can show how increasing the number of replicates increases the probability of detecting a given level of isotope incorporation. Power analysis can also show, at constant level of power, how increasing the number of replicates decreases the threshold level of isotope incorporation that can be detected. Lastly, power analysis can clarify the tradeoffs between Type I and Type II errors, which can provide useful context for interpreting results from qSIP experiments.

One way to address issues inherent to metagenomic analysis (e.g. higher amounts of DNA required for sequencing, higher sequencing costs and exponentially increased computational complexity) is to reduce the number of density fractions. In addition, adding replication with a reduced number of fractions (gradient resolution) could lead to higher accuracy while maintaining a similar effort to high gradient resolution without replication. We investigated the repercussions of reducing the number of density fractions on replicated and unreplicated datasets from marine and terrestrial microbial communities using different isotopes.

Using multiple SIP datasets, we tested the robustness of qSIP with variation in density fraction size. We combined (in-silico) density fractions from real datasets and measured the effects of lower gradient resolution on per-taxon density shifts and unlabeled weighted mean density. We show that reducing the gradient resolution from an average density fraction size of 0.002 g ml^-1^ down to 0.011 g ml^-1^ (50 to 9 fractions of a 5 ml tube) yields comparable shift detection with a detection limit of 0.005 g ml^-1^ (9% enrichment with ^13^C). We discuss using the small inherent variability between replicates as a way to define a shift detection limit. Finally, we show that this inherent variability is more similar between replicates centrifuged together (within spin) than between replicates centrifuged separately (between spins), stressing the need for internal standards that can be spiked into each sample rather than external standards.

## Methods

We used five datasets representing different ecosystems for *in silico* analyses: a high resolution unreplicated SIP study of ^13^naphthalene in seawater, two medium resolution replicated SIP experiments where ^18^O-water was added to soils, replicated genomic DNA from pure cultures of *Escherichia coli* and *Pseudomonas putida*, and a replicated genomic mock community comprised of high molecular weight genomic DNA of *Thermoanaerobacter pseudethanolicus, Bacillus licheniformis, Bifidobacterium longum* subsp. Infantis and *Streptomyces violaceoruber* purchased from ATCC. See table 1 for number of density fractions and number of replicates per dataset.

**Table 1:** Datasets used in this study including source, number of replicates and analyses performed

| Dataset | Replicates | Fractions |
| --- | --- | --- |
| Naphthalene enriched seawater | 1 | 50 |
| SPRUCE peatland | 5 | 18 |
| Grassland | 3 | 9 |
| *E. coli* and *P. putida* | 4-6 | 30 |
| Mock community | 9 | 48 |

As these experiments were performed by different laboratories, using slightly different protocols, we describe their SIP pipelines separately. However, all post-sequencing steps were performed identically for all 16S-rRNA operational taxonomic units (OTUs).

### Naphthalene enriched seawater dataset

Surface seawater was collected from the port of Los Angeles in July 2014 and May 2015. Ten liters of water were incubated at ambient temperature with 400 nM ^12^C- or ^13^C-naphthalene for 24 hours (July 2014) or 88 hours (May 2015). The water was then filtered sequentially through an 80 nM mesh, a 1 µ prefilter (Acrodisc syringe glass fiber, Pall Laboratory) and a 0.2 µ polyethersulfone (PES) Sterivex filter (Millipore). After filtration, 1.5ml Sodium-Chloride-Tris-EDTA (STE) buffer was injected into the Sterivex casing and the filters were promptly sealed and stored in −80°C.

DNA was extracted from the Sterivex filters by bead beating (10 minutes), transferring the lysate into DNeasy plant kit columns (Qiagen) and following the kit protocol. The eluted DNA was stored in −80°C until use.

In preparation for ultracentrifugation, 1 µg of eluted DNA from the labeled (^13^C) and control (^12^C) incubations was mixed with CsCl solution and gradient buffer (GB) for a final density of 1.725 g ml^-1^. The gradients were centrifuged in a Beckman NVT 65.2 rotor at 44,100 rpm for 64-68 h at 20°C. Following centrifugation each gradient was manually fractionated into 50 equal fractions of 100 µl each. The density of each fraction was determined using a handheld AR200 digital refractometer by removing 10 µl per fraction.

DNA in each fraction was purified and concentrated using glycogen/PEG precipitations followed by an ethanol washing and elution in Tris-EDTA buffer (TE). DNA was then quantified by PicoGreen assay (Life Technologies). The 16S-rRNA coding gene hypervariable regions V4-V5 were amplified from each fraction that contained DNA using universal primers 515F-N and 926R (Parada, Needham, and Fuhrman 2016). Each reaction tube contained 10 µM of each primer, 1 ng of DNA, 12 µl 5Prime Hot Master Mix and 10 µl PCR water. The thermocycler was set to 3 minutes denaturation at 95°C; 30 cycles of: denaturation 95°C 45 seconds, annealing 50°C 45 seconds and elongation 68°C 90 seconds; followed by a final elongation step at 68°C for 5 minutes. PCR products were cleaned using 1x Agencourt AMPure XP beads (Beckman Coulter), quantified via PicoGreen, and pooled in equimolar amounts and sequenced on Illumina MiSeq 2×300bp. Mock communities and PCR blanks were included in all sequencing runs (Yeh et al. 2018).

### Soil ^18^O-water dataset 1: spruce peatland

Soil samples (0-10cm; n=5) were collected from the Marcell Experimental Forest, located in northern Minnesota in August 2017. Samples were then air-dried in the lab to minimize O16-water content. One-half gram dry weight soil was weighed into 15mL Falcon tubes and pre-incubated at one of five temperatures (n=20) for approximately 48 hours in the dark: 5C, 15C, 25C, or 35C. After pre-incubation, half the samples received enough natural abundance O16-labeled water (n=10) to bring the sample up to 60% field capacity, and the other half (n=10) received 97-atom % O18-labeled water. Samples were placed back at their original incubation temperature and harvested after 5 (n=5) and 10 days (n=5). Lids of the Falcon tubes were opened every ∼24 hours to allow for CO2 release. Samples were frozen at −80C until further processing. DNA was extracted using a PowerSoil DNA extraction kit following manufacturer’s instructions (MoBio Laboratories, Carlsbad, CA). For stable isotope probing, approximately 1 g of DNA was loaded into a 4.7 mL ultracentrifuge tube with 6.88 g of a saturated cesium chloride solution and filled with gradient buffer (200 mM Tris, 200 mM KCl, 2 mM EDTA). Samples were spun in a Beckman OptimaMax benchtop ultracentrifuge (Indianapolis, IN, USA) using a Beckman TLA-110 rotor at 150,200 x g at 18C for 72 hours. Tubes were fractionated into approx. 20 fractions of 200 L each and the density of each fraction was measured with a Reichart AR200 digital refractometer (Buffalo, NY, USA). DNA was purified using a standard isopropanol precipitation method and quantified by PicoGreenTM fluorescence on a BioTek Synergy HT plate reader (Winooski, VT, USA).

The V3-V4 region of the 16S rRNA gene was subsequently quantified and sequenced in samples within the density range of 1.640 – 1.746 g mL-1 (approx. 15 fractions per sample). To quantify the 16S rRNA gene, qPCR was performed in triplicate using a Bio-Rad CFX384 Touch real-time PCR detection system (Hercules, CA, USA) and primers Eub338F (5’-ACTCCTACGGGAGGCAGCAG-3’) and Eub518R (5’-ATTACCGCGGCTGCTGG-3’) (Fierer et al. 2005). The 10 µL qPCR reactions contained 1 µL of sample and 9 µL of master mix (0.25 mM of each primer, 1X Forget-Me-Not EvaGreen qPCR mix (Biotium, Fremont, CA), and 0.4 mg mL-1 BSA. The PCR program used was as follows: 95°C for 2 min, followed by 40 cycles of 95°C for 30s, 59°C for 10s, and 72°C for 10 sec.

For sequencing, two PCR steps were used to process the samples, as in Berry et al. (2011). Each sample was first amplified using primers 515F (Parada) (5’-GTGYCAGCMGCCGCGGTAA-3’) and 806R (Apprill) (5’-GGACTACNVGGGTWTCTAAT -3’) (Apprill et al. 2015; Parada, Needham, and Fuhrman 2016). This was done in duplicate 10 µL PCR reactions containing 1 µL of DNA template and 9 µL of master mix (1 µM of each primer, 1X Phusion Green HotStart II Polymerase (Thermo Fisher Scientific, Waltham MA) and 1.5 mM MgCL2). PCR conditions were 95°C for 2 min, then 15 cycles of 95°C for 30s, 55°C for 30s and 72°C for 10 s. Initial duplicate PCR products were pooled, checked on a 1% agarose gel, 2-fold diluted, and used as template in the subsequent tailing reaction with the same primers that included the Illumina flowcell adapter sequences and a 12 nucleotide Golay barcode (15 cycles identical to initial amplification conditions). Amplicons were then purified with 0.1% carboxyl-modified Sera-Mag magnetic Speed-beads (Thermo Fisher Scientific, Freemont, CA, USA) in 18% PEG and quantified with a PicoGreenTM assay on a BioTek Synergy HT plate reader. Samples were then pooled at the same concentration, purified again with the Sera-Mag beads as described above, and quantified with the KAPA Sybr Fast qPCR kit (Wilmington, MA, USA). Libraries were sequenced on an Illumina MiSeq (San Diego, CA, USA) instrument at Northern Arizona University’s Environmental Genetics and Genomics Laboratory, using a 300 cycle v2 reagent kit.

### Soil 18O-water dataset 2: Grassland

A 10 x 4.5 cm soil core was collected using an AMS 15 x 4.5 cm soil core sampler from the upper layer of soil at the Buck field site at Hopland Research and Extension Center, Hopland California in February 2018, and transported on wet ice and stored at 4C for one month. The core was homogenized and split into two microcosms; half of the microcosms were wetted with ^16^O-H_2_O and the other half with 97-atom% ^18^O-H_2_O and incubated for 8 days at room temperature. DNA was extracted from each sample using the PowerSoil DNA extraction kit following manufacturer’s instructions (MoBio Laboratories, Carlsbad, CA). DNA was added to cesium CsCl solution and gradient buffer (GB) for a final density of 1.725 g ml^-1^. The gradients were centrifuged in a Beckman VTi 65.2 rotor at 44,100 rpm for 109 h at 20°C. Following centrifugation each gradient was fractionated into 38 equal fractions of 135 µl each. The density of each fraction was determined using a handheld AR200 digital refractometer by removing 5 µl per fraction. DNA in each fraction was purified and concentrated using glycogen/PEG precipitations followed by an ethanol washing and elution in Tris-EDTA buffer (TE). DNA was then quantified by PicoGreen assay (Life Technologies). Fractions were pooled to nine sets, encompassing 1.6900-1.7099, 1.7100-1.7149, 1.7150-1.7199, 1.7200-1.7249, 1.7250-1.7299, 1.7300-1.7349, 1.7350-1.7399, 1.7400-1.7468, and 1.7469-1.7720 g/mL density ranges. DNA from each fraction—as well as unfractionated DNA from each mesocosm—was fragmented using the Bioruptor Pico sonicator (Diagenode Inc, Denville, NJ) and prepared for shotgun metagenomic sequencing using the Kapa Hyperprep Plus kit with 3 rounds of PCR amplification (Kapa Biosystems, Wilmington, MA), and sequenced to an average depth of 7 gbp per fraction on the Illumina NovaSeq (Illumina, San Diego, CA).

Reads from each library were processed for PhiX and adapter contamination using bbduk (https://jgi.doe.gov/data-and-tools/bbtools/bb-tools-user-guide/bbduk-guide/) and low-quality base pairs trimmed using sickle (https://github.com/ucdavis-bioinformatics/sickle) with default settings. Trimmed reads for all fractions from each mesocosm were assembled together with megahit (Li et al. 2015) with --k-min 21 --k-step 6 --k-max 255. Reads from each fraction from both mesocosms were mapped to each metagenomic assembly using bbmap (https://jgi.doe.gov/data-and-tools/bbtools/bb-tools-user-guide/bbmap-guide/) with fast=t ambig=random and minid=0.98. Metagenomic contigs from each assembly were binned into draft microbial genomes using metabat2 (Kang et al. 2019). Reads from trimmed libraries were mapped again to one genome bin of interest with bowtie2 (Langmead and Salzberg 2013), read-depth calculated in 1kb windows across the genome bin using bedtools coverage (Quinlan 2014), and visualized with custom r scripts relative to average GC (calculated using custom python scripts).

### Genomic mock communities and pure cultures

DNA for the genomic mock communities was purchased from ATCC, resuspended in Tris-eDTA buffer, mixed in equal proportions and aliquoted into replicates. The mock communities were composed of high molecular weight DNA of *Thermoanaerobacter pseudethanolicus, Bacillus licheniformis, Bifidobacterium longum* subsp. Infantis and *Streptomyces violaceoruber* (see sup. Table S1 for accession numbers). These genomes were selected for their distinguishable %GC content (34.5%, 46%, 60% and 73% respectively). These mock communities as well as DNA extracted from pure cultures of *Escherichia coli K-12* and *Pseudomonas putida KT2440* was centrifuged in a Beckman VTi 66.2 rotor at 20°C for 120 hours at 44,000 RPM. These samples were fractionated by Agilent 1260 Infinity II analytical-scale fraction collector with isocratic pump followed by precipitation, washing and elution by a Hamilton Vantage pipetting robot. DNA was quantified using Quant-iT DNA High Sensitivity Assay.

### 16S-based microbial community composition in individual fractions

Each dataset was processed separately. Reads were quality-trimmed using Trimmomatic version 0.33 (Bolger, Lohse, and Usadel 2014) with parameters set to LEADING:20 TRAILING:20 SLIDINGWINDOW:15:25. The resulting reads were merged using Usearch version 7 (Edgar 2010), clustered in Mothur following the MiSeq SOP and classified using the Silva taxonomy database version 119 (Schloss et al. 2009; Pruesse, Peplies, and Glöckner 2012; Kozich et al. 2013).

To track individual operational taxonomic units (OTUs) over density fractions, the relative abundance of the OTU in each fraction was multiplied by either the concentration of DNA in the same fraction (seawater) or the total 16S copy number (soil dataset 1). The results were normalized to the total abundance of that OTU over all fractions for an area of 100% under each curve.

### Density shifts

The weighted mean density of each OTU in labeled and control samples was calculated by multiplying density by OTU abundance (amount of DNA/16S copies * OTU relative abundance) within each fraction, summing up the products and dividing them by the sum of abundances of the OTU across all fractions. The weighted mean density shift was calculated by subtracting the weighted mean density of the OTU in the natural abundance treatment from the labeled sample. The density shifts were plotted in R (Team 2018) for the 100 most abundant OTUs in each sample.

Relative error was calculated as: let r be a gradient resolution (r < original number of fractions (r_max_)). The relative error is the difference between the density shift per OTU in resolution r minus the shift per OTU in r_max_.

### Sensitivity analysis

Using unlabeled replicated (N=10) samples from soil dataset 1, we calculated the weighted mean density and its standard deviation for each of the 100 most abundant OTUs. We used two standard deviations as the detection limit per OTU under the assumption that a shift that is smaller than or equal to the natural variability in the unlabeled weighted mean is not detectable. For the OTU abundance effect on WMD variability we used 320 OTUs from the same dataset.

### Sensitivity to number of fractions

We used datasets 1 and 2 to estimate how precision in the estimate of isotope incorporation varies with the number of density fractions collected in a qSIP experiment. We combined fractions in silico to simulate experiments with fewer fractions (f), with the following principles: 1) Only adjacent fractions were combined. 2) Fraction combinations were conducted in order to create new, combined fractions that were approximately equal in size and sequencing depth (i.e., with minimal variation in the range of densities represented by each simulated fraction). For example, to simulate an experiment where only two density fractions were collected, we ran three possible scenarios: combining the lightest 8, 9 or 10 fractions into one, simulated fraction and combining the heaviest 8, 9, or 10 fractions into a second, simulated fraction (9 v 9, 8 v 10, or 10 v 8). We did this to simulate typical approaches to SIP experiments, where fractions that span similar density ranges are typically selected. For each permuted combination in the replicated dataset 2, we ran the qSIP code (https://rdrr.io/github/bramstone/qSIP/f/README.md) and estimated atom percent excess ^18^O for each replicate tube, and then calculated the standard deviation in that estimate across all replicates (n=5). Finally, we calculated the relative standard deviation as a function of increasing number of fractions included in the simulation compared to the original number of fractions.

### Power analysis

We evaluated statistical power using the SPRUCE dataset. We used data from soils incubated for 10 days at an intermediate temperature (15 °C), sampled after 5 and 10 days of exposure to ^18^O-H_2_O. The unlabeled control was sampled at day 0. The power analysis focused on taxa that occurred in all 15 samples (n=5 for control, ^18^O-H_2_O at day 5, and ^18^O-H_2_O at day 10), omitting taxa in the uppermost 5^th^ percentile for standard deviation of the estimate of weighted average density, which are likely to be rare taxa (see Figure 1). We used observed variation among taxa for day 5 and day 10 in weighted average density shift, which ranged from −0.003 to 0.033 g cm^-3^. This captures a wide range of possible values of isotope uptake, from ∼0 to ∼60 atom percent excess ^18^O. We used resampling with replacement to estimate power. For each taxon at each sample date, *N* random samples were drawn (with replacement) from each the ^18^O-labeled and unlabeled datasets, a t-test was performed, and the P-value was recorded. This was repeated 1000 times, and power was estimated as the frequency of significant t-tests among the 1000 simulations. N was varied to simulate experiments with different numbers of replicates by pruning or duplicating replicates from the original dataset, ranging from N=2 to N=6. Average power was calculated across all taxa. The upper 10^th^ percentile was also calculated to estimate power typical for more dominant taxa.

**Figure 1:**
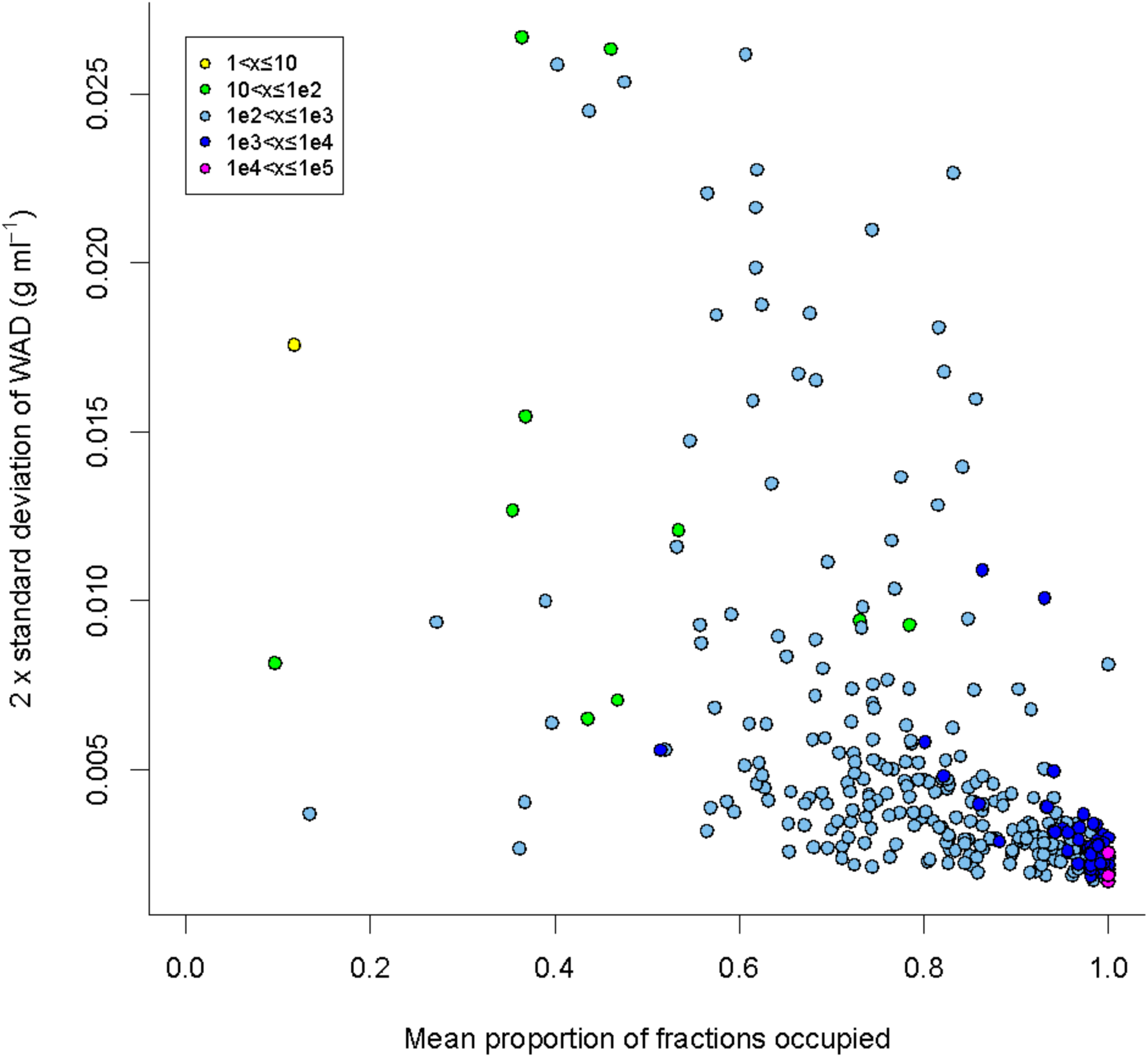
The effect of OTU abundance on qSIP sensitivity. OTU abundance is positively correlated to the number of fractions in which it can be detected and negatively correlated to the standard deviation of its unlabeled weighted mean density. Coefficient of variation (standard deviation divided by the mean) of the weighted mean density of 320 from soil dataset 1. As a function of the proportion of density fractions the OTU appears in. The colors represent the abundance of the OTU in the unfractionated sample in 16S copy number.

## Results

### Abundance is negatively correlated to qSIP variation

Density shifts, or change in weighted mean density (WMD), due to incorporation of a stable isotope labeled substrate are the basis for calculating isotope enrichment. Those shifts have been shown in-silico to be detectable with qSIP in moderately to highly abundant OTUs (>0.1% relative abundance) (Youngblut, Barnett, and Buckley 2018). First, we set out to ground-truth this finding using experimental data. We show that the variability of the unlabeled WMD is negatively correlated to the abundance of OTUs. Namely, the more abundant an OTU is -the more consistent its WMD is (Fig. 1).

The physics of the behavior of DNA within density gradient centrifugation affects the number of fractions in which presence of OTUs can be detected. The long tails of DNA to density distributions are attributed to a smear of DNA along the tube wall (Youngblut, Barnett, and Buckley 2018). It stands to reason that the more abundant an OTU - the higher its representation in this smear will be. In addition, the detection limit of an OTU affects the number of fractions it will be detected in. Indeed, we show that when inspecting presence/absence of OTUs in all fractions, OTU abundance is positively correlated with the proportion of fractions in which it is present (Fig. 1). Abundant OTUs appear in almost all fractions, whereas rare OTUs appear in few fractions, and in some cases only one fraction. These rare OTUs also have highly variable WMD values.

### Density shifts are consistent across medium to high gradient fractionation resolution

We started with an unreplicated dataset of OTUs from naphthalene-enriched seawater DNA divided into 50 fractions of which 45 had quantifiable DNA. Consecutive density fractions were consolidated *in-silico* (every 2-, 3-, 4 fractions etc) to represent a range of fraction sizes spanning 0.002-0.02 g ml^-1^, and the density shift of the 100 OTUs that were most abundant in all fractions combined was examined. The estimated magnitude of the density shifts across taxa remained consistent at a fraction size of up to 0.011 g ml^-1^, expanding previous results that demonstrated this trend with fractions of 0.003-0.008 g ml^-1^ (fig. 2A) (Youngblut, Barnett, and Buckley 2018). The same data can be represented as a relative error, which is defined here as the density shift in resolution r (r < original number of fractions) compared to the density shift with maximum resolution. When the relative error is high there is a higher probability of mis-assigning taxa as incorporators when they are not and vice versa. There was a positive linear correlation (R^2^ = 0.95) between fraction size and mean relative error (fig. 2B). Additionally, the increase in the mean relative error is accompanied by an increase in its variation, further emphasizing the risk of type II errors.

**Figure 2:**
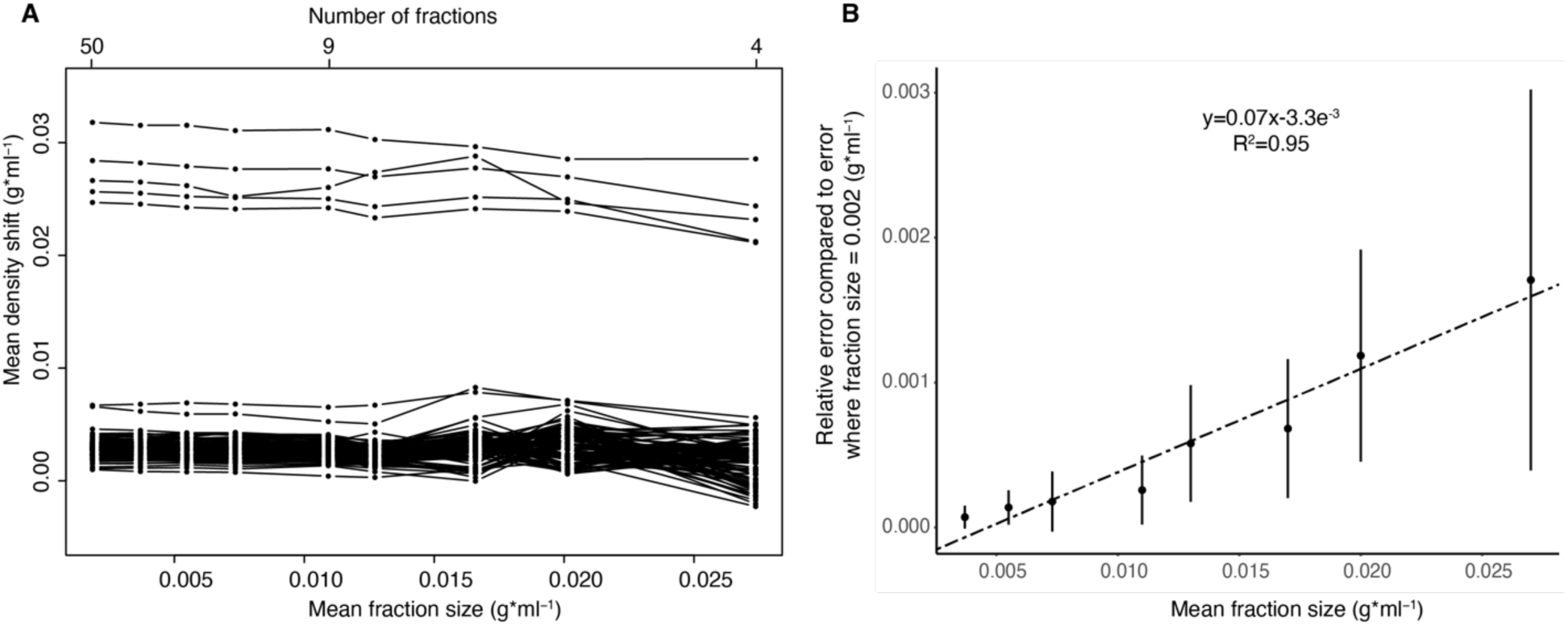
variation in estimated isotopic enrichment declines with smaller density fractions in unreplicated data. **(A)** Mean fraction size, while smaller than 0.011 g ml^-1^ (corresponding to 9 fractions in a 5 ml ultracentrifuge tube), does not affect the density shift of OTUs. Each line represents one OTU. The plot shows the density shift (y axis) of the top 100 most abundant OTUs in seawater enriched with naphthalene. Highly enriched taxa are easily discerned even at a fraction size of 0.02 g ml^-1^ (4 fractions), but shifts of less or not enriched OTUS, while remaining within a narrow range, may increase or decrease and negatively affect % atom excess downstream analyses. **(B)** Relative error was calculated as the absolute difference between density shift in a fraction size and the density shift when the mean fraction size was 0.0018 g ml^-1^ per OTU from the same data. RelErr = Mean(Shift_r_ - Shift_max_) where r is a gradient resolution lower than the maximum.

However, when adding replicates, the correlation between the number of fractions on the relative standard deviation of the WMD is no longer linear. Between 2 and 11 fractions, every additional fraction reduces the standard deviation exponentially, whereas with at least 12 fractions the difference is much smaller and linearly correlated to the number of fractions (fig. 3B).

**Figure 3:**
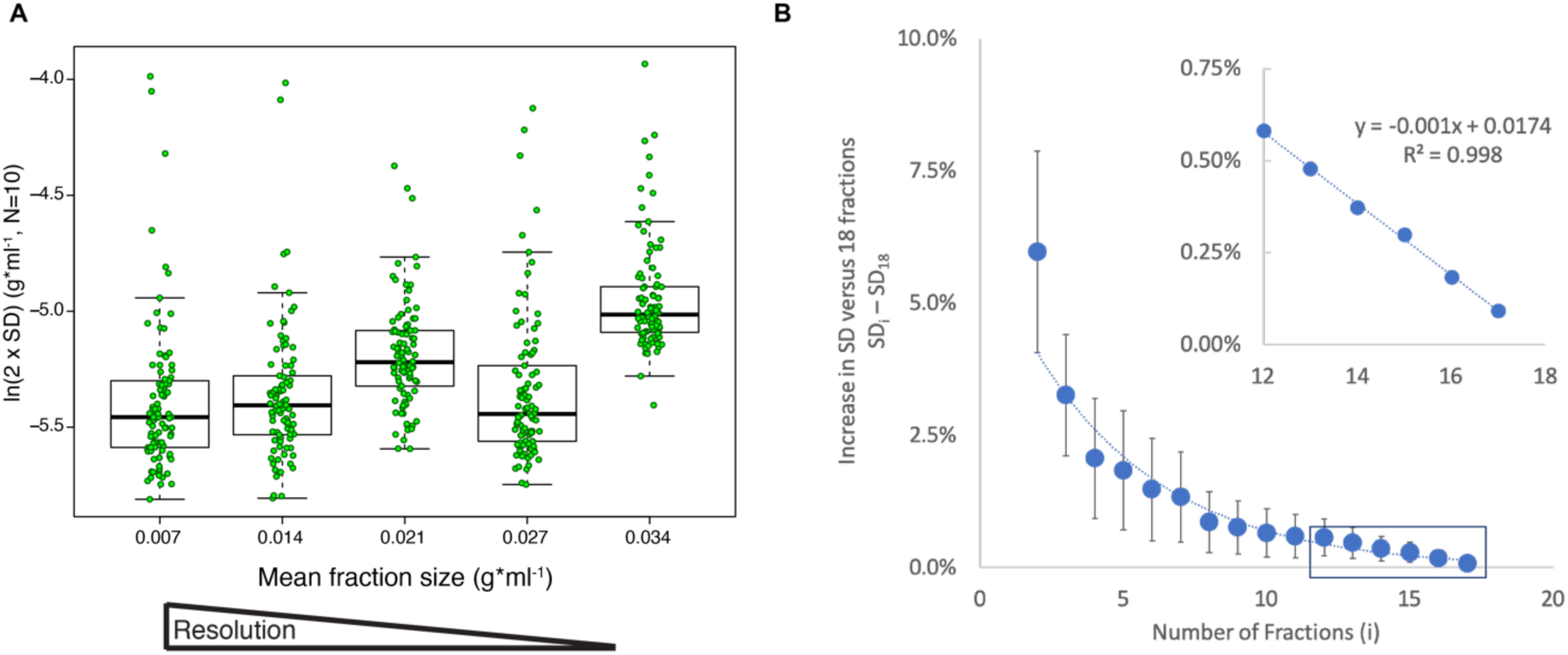
Variability in level of enrichment is negatively correlated to the number of density fractions in replicated data. **(A)** Two standard deviations of the mean buoyant density in the 100 most abundant OTUs from the unlabeled terrestrial dataset (N=10) as a function of fraction size. Higher fraction size corresponds to lower gradient resolution. The horizontal line within the box represents the median, the box represents percentiles 25-75 and whiskers represent percentiles 10 and 90 for 100 OTUs in each fraction size. The raw data is plotted on top of the boxes. **(B)** Relative error compared to the original 18 fractions dataset (N=5, 100 permutations) decreases as the number of fractions increases. The inset shows a linear decrease in relative error when using 12-18 fractions.

### Low inherent variability determines density shift detection limit

To identify statistically significant density shifts between samples treated with a labeled substrate versus a control (unlabeled substrate), it is critical to know the detection limit of density shifts. To define a detection limit, we calculated the inherent variability in weighted mean density of unlabeled DNA from various taxa in a highly replicated experiment enriching soil with 18O-water (N=10). We extended this analysis to explore the impact of gradient resolution reduction on this variability. This was done by merging and averaging data from an increasing number of consecutive fractions. The initial analysis with medium resolution (fraction size 0.007 g ml^-1^; 11-17 fractions in a 4.7 ml tube) revealed that the weighted mean density of abundant taxa varied little between replicates (fig. 3A). At a 95% confidence level (two standard deviations), the mean of replicates per taxon varied at a median value of 0.004-0.007 g ml^-1^ at gradient resolutions varying from 0.007-0.034 g ml^-1^ variation of 90% of the taxa was always lower than the fraction size. The variation remains comparable with lower gradient resolution down to a fraction size of 0.027 g ml^-1^, and only increases significantly (ANOVA, Tukey 95% confidence level) at a fraction size of 0.034 g ml^-1^ (3 fractions in a 4.7 ml tube) (fig. 3A). Further analysis of the increase in standard deviation compared to the standard deviation with the original 18 fractions revealed a linear increase between 12 and 18 fractions, and an exponential increase with 2-11 fractions (fig. 3B).

Additionally, the range of relative error increased with fraction size. To explore how this variation affects the detection of substrate incorporators, we calculated the putative sensitivity (proportion of true positives) and specificity (proportion of true negatives) as a function of the shift detection threshold for all gradient resolutions discussed previously. This calculation was performed under the assumption that a density shift higher than a specific threshold in the original experimental setup (50 fractions) represented significant enrichment. The shift detection threshold is the smallest difference between labeled and unlabeled WMD that would be considered a significant density shift. As expected, both parameters were stable down to 0.011 g ml^-1^ density fraction resolution using a shift detection threshold 0.005 g ml^-1^ or higher. Specificity was > 95% for all gradient resolutions at a threshold > 0.003 g ml^-1^, but sensitivity was more impacted by gradient resolution > 0.013 g ml^-1^ (fig. 4).

**Figure 4:**
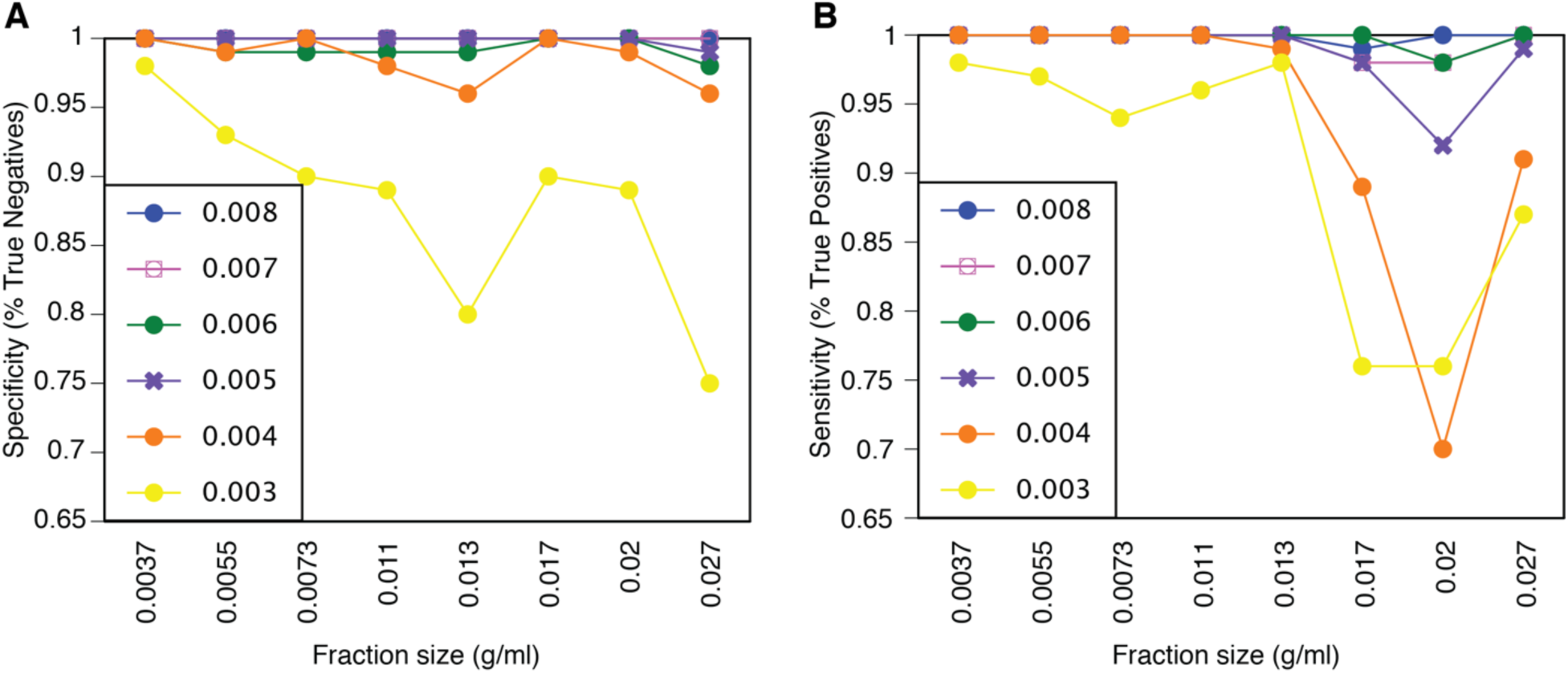
Rate of false discoveries increases at low gradient resolution or low shift detection threshold. Specificity (A; rate of true positives) and sensitivity (B; rate of true negatives) calculated over the 100 most abundant OTUs from naphthalene-enriched seawater. The colors represent detection limit thresholds.

In a replicated experiment, the number of replicates and desired statistical power determine the detection limit. When designing an experiment, it could be valuable to use the desired statistical power and desired enrichment detection threshold to decide on the number of replicates. Both of these parameters would depend on the scientific purpose of the study. The higher the power and threshold, the less replicates are necessary (fig. 5).

**Figure 5:**
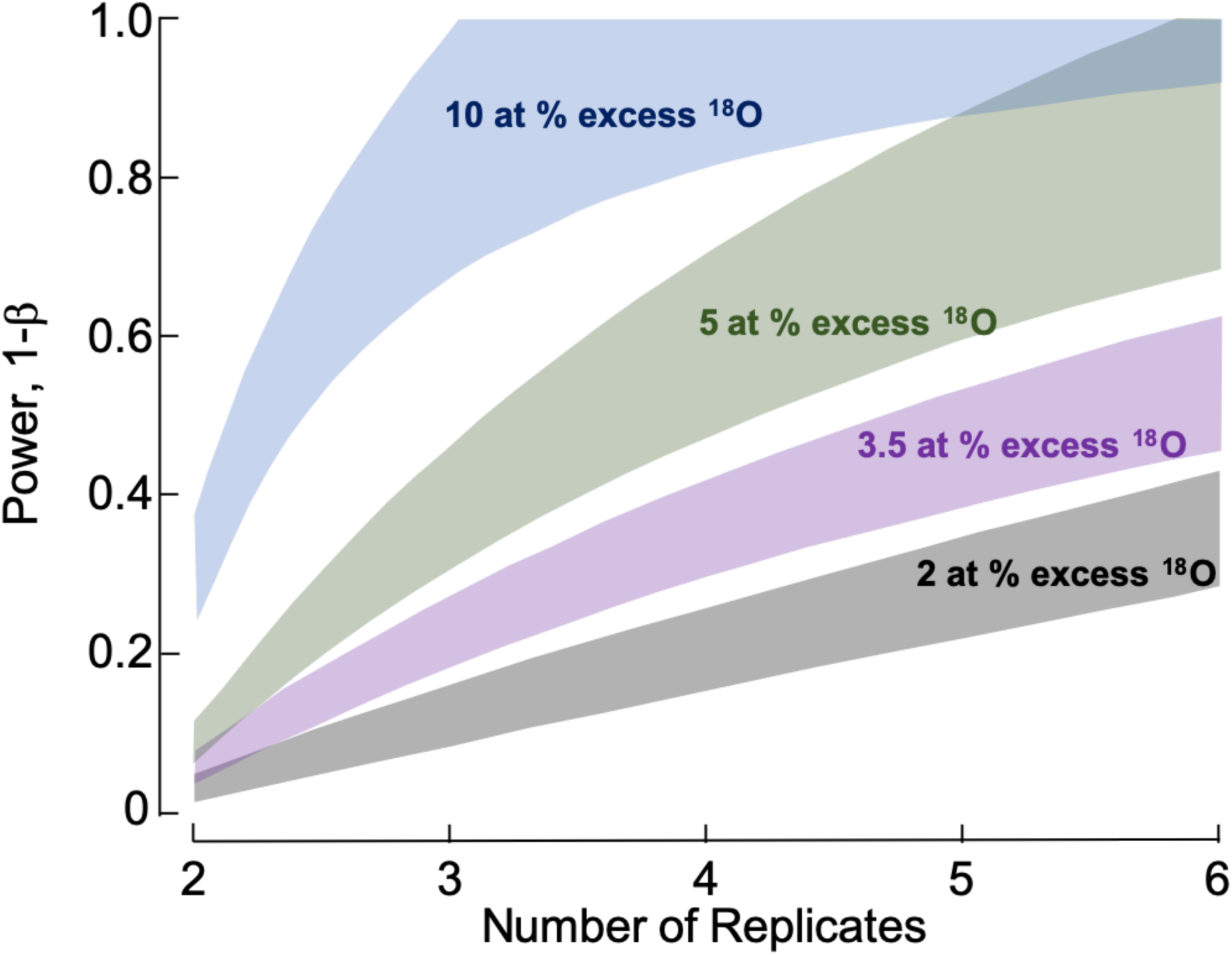
The number of replicates of a qSIP experiment determines the statistical power of enrichment detection and the detection limit. 10 atom percent enrichment (APE) by incorporation of 0.0065 g ml^-1 18^O labeled substrates would correspond to 12.6 APE with the same incorporation of ^13^C substrates or 6.3 APE with ^15^N substrates.

### Variation in mean weighted density is greater within than between spins

DNA extracted from pure cultures of unlabeled *Escherichia coli* and unlabeled or 100% ^13^C-labeled *Pseudomonas putida* was aliquoted into replicates, ultracentrifuged in CsCl and fractionated. The difference in %GC of the genomes of these organisms (i.e. distance between their peak densities) permitted calculation of their WMD even when both were unlabeled. Comparing the mean of WMD of replicates of E. coli between spins revealed negligible variation. However, the range of mean weighted densities within a spin was up to 0.004 g ml^-1^ (fig. 6).

**Figure 6:**
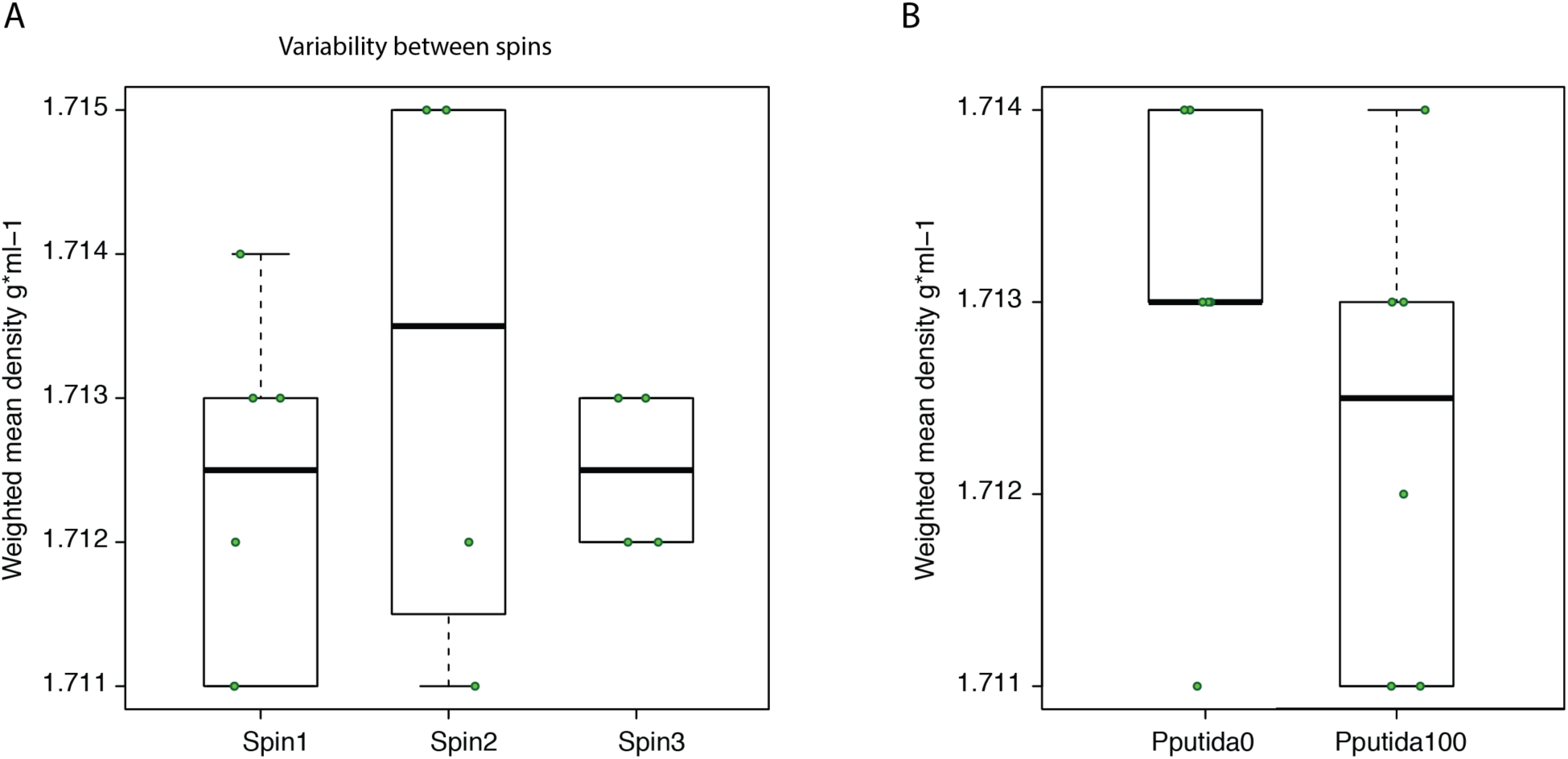
Tube-to-tube variability and effect of other taxa. Variability in the weighted mean density (rounded to 3 decimal places) of unlabeled *Escherichia coli* (A) between spins (N=6, N=4, N=4 respectively) in the presence of labeled *Pseudomonas putida* at 100 atom% ^13^C and (B) within the same spin in the presence of unlabeled vs. labeled *P. putida* at 100 atom% ^13^C (N=6).

Similar results were obtained by spinning triplicates of genomic mock communities comprised of four organisms with distinguishable genomic %GC. In these experiments between-spin variation was 0.0013-0.0025 g ml^-1^, whereas within-spin variation ranged up to 0.0056 g ml^-1^ (sup. Fig. S1, sup. table S2 (raw data)).

### Using genomic mock communities to explore density to GC content conversion

One potential strategy for decreasing the cost of metagenomic SIP experiments is to sequence only the labeled samples and calculate an approximate density shift using the %GC of the genomic bins. The conversion of density to the GC content of a genome is a linear function of the unlabeled (^12^C) weighted mean. There is a canonical equation describing this function (Rickwood and Birnie 1978; Schildkraut, Marmur, and Doty 1962; Buckley et al. 2007) but it has also been determined empirically in the past by using a ladder of three bacterial taxa with varying GC contents (Hungate et al. 2015). However, if the equation was identical for each gradient in a SIP experiment, as is usually assumed, unlabeled replicates of the same organism would have had the exact same weighted mean in every run. As shown here and previously, this is not the case (Hungate et al. 2015; Morando and Capone 2016). To address this variation, we ran 9 replicates of a mock community with a wide range of known GC content, generated a calibration curve from each one and fitted an equation to it.

This mock community, consisting of high molecular weight genomic DNA from 4 bacterial taxa, revealed highly correlated linear relationships (N=9, R^2^>0.94) between mean weighted density and %GC, but with a variance in slopes and intercepts (table 1). There was a significant difference between the weighted mean density as calculated according to the canonical equation and the observed mean of weighted mean density per genome over all replicates (N=9, paired t-test, p=0.02). For example, using the canonical equation (Schildkraut, Marmur, and Doty 1962) on replicate 1 would lead to GC content of the mock community members to appear as 38%, 46.5%, 62% and 72.1% respectively. We also noticed that the difference between observed and expected mean weighted density decreased as %GC increased (table. 2).

**Table 2:** Observed vs. expected GC content for known genomes varies between replicates. Calculated %GC (Schildkraut, Marmur, and Doty 1962), slope and intercept between replicates of the genomic mock community. The header row shows the known %GC per genome.

| Replicate | 34.5% | 45.9% | 60.1% | 72.7% | slope | intercept | R<sup>2</sup> |
| --- | --- | --- | --- | --- | --- | --- | --- |
| 1 | 37.8% | 46.9% | 62.2% | 72.4% | 0.0904 | 1.6654 | 0.995 |
| 2 | 37.8% | 52% | 64.3% | N/A | 0.0985 | 1.6642 | 0.984 |
| 3 | 40.8% | 48% | 64.3% | 75.5% | 0.0934 | 1.6661 | 0.993 |
| 4 | 39.8% | 48% | 64.3% | 76.5% | 0.0978 | 1.6639 | 0.995 |
| 5 | 40.8% | 50% | 66.3% | 78.6% | 0.0989 | 1.6652 | 0.997 |
| 6 | NA | 45.9% | 53.1% | 68.4% | 0.0817 | 1.666 | 0.943 |
| 7 | 39.8% | 50% | 63.3% | 74.5% | 0.0904 | 1.6677 | 0.999 |
| 8 | 40.8% | 51% | 63.3% | 75.5% | 0.0889 | 1.6689 | 0.9998 |
| 9 | 42.9% | 52% | 65.3% | N/A | 0.0869 | 1.6713 | 0.9998 |
| mean | 39.8% | 49% | 63.3% | 74.5% | 0.0895 | 1.6678 | 0.9998 |

## Discussion

SIP is a powerful tool for investigating taxon-specific microbial functions in complex assemblages. Like any method, SIP-derived measurements have some inherent variability which can be managed – within limits – to address the research questions of interest. Our results show how this can be done given a particular research question and the level of sensitivity / detection demanded by that research question. Despite the wide use of SIP, there has been little benchmarking of interpretation of its results. Here we attempted to shed light on some practical aspects of the method and discuss how to adjust it to maximize results.

Variation of the mean weighted density (WMD) of the same unlabeled taxon over numerous replicates, observed even between samples processed simultaneously and in the same manner, implies that there are unpredictable physical and/or chemical factors unrelated to genomic %GC affecting SIP analyses. The variability these factors create can determine the limit of density shift detection. Both the unlabeled WMD variation and interpretation error analyses performed here imply that when using qSIP a detection limit balancing Type-I and Type-II errors may revolve around 0.005 g ml^-1^ in unreplicated ^13^C experiments when dividing the density gradient into at least 4 density fractions. Replication would lead to increased statistical power using the same detection limit. However, density-shift estimates of taxon-specific isotope incorporation are broadly robust across a wide range of fraction sizes. For example, the relative error of high incorporators only varied by an average of 0.02% in shift from fraction size 0.002-0.011 g ml-1 (N=5, 50 to 9 fractions in a 5.1 ml tube).

In reality, WMD variation means that the same organism may peak at a density anywhere within a specific range. Moreover, the unlabeled mean and labeled mean of a single replicate can deviate in different directions, increasing the observed density shift and leading to a Type I error, and the potential for such deviation increases at low gradient resolution. This may explain why the variation in the relative error increases as resolution decreases.

Low gradient resolution in combination with higher variability may also lead to false classification of borderline taxa as incorporators when they are not (Type I error) and vice versa (Type II error). For example, a simulation model (Youngblut, Barnett, and Buckley 2018) showed that the rate of true negatives (specificity) and true positives (sensitivity) of qSIP is 88% and >90% respectively, with virtually no effect of fraction size at the range of 0.003-0.008 g ml-1. When examining the specificity and sensitivity of qSIP using real unreplicated data over a wider range of fraction sizes we found that gradient resolution and shift detection limit both had an effect. However, the reliability of qSIP remained extremely high as long as the detection limit was 0.005 g ml^-1^ or higher (Specificity > 90% and sensitivity > 95%) regardless of gradient resolution. This detection limit is comparable to 2 standard deviations of unreplicated unlabeled weighted mean density, further supporting that analysis. Experimental replication, even as low as 3 replicates, can increase the power of this analysis to have virtually no error using a similar detection threshold. Our analysis can be used for experimental design based on the desired statistical power.

The reduction of the number of density fractions we addressed in this study could significantly mitigate labor and sequencing costs. While helpful for amplicon-SIP, this reduction is crucial for metagenomic SIP (MG-SIP). The combination of metagenomics and SIP, first attempted over a decade ago (Dumont et al. 2006; Schwarz, Waschkowitz, and Daniel 2006), involves sequencing metagenomes instead of amplified marker genes from density fractions, and assembly of genomic bins from those metagenomes. Genomic bins that shift to a higher density can then be identified and their metabolism explored directly. Many of the obstacles that were brought up in the past with regards to MG-SIP (Chen and Murrell 2010) were addressed by the improvement in sequencing platforms and library preparation kits, such as low DNA yield, MDA biases and low throughput. Still, so far studies combining metagenomics and SIP included shotgun sequencing of labeled DNA, a few heavy fractions, or at best also sequenced 2-3 light fractions (Dombrowski et al. 2016; Fortunato and Huber 2016; Thomas, Corre, and Cébron 2019). This approach limits detection of substrate incorporators in several ways: (1) choosing which fractions to sequence relies on density shift of the entire community, which may be subtle (2) low GC genomes, even if highly enriched, may not become heavy enough to reach the heavy fractions, (3) low GC genomic islands may not be well-covered and therefore not assembled for the same reason, leading to increased genome fragmentation (sup. fig. S2), (4) depending on the density of the fractions sequenced, high GC genomes may be highly represented regardless of enrichment, (5) an organism may be abundant in the sample but not be enriched enough to reach the heavy fractions due to additional use of other substrates, in which case it could take up a good amount of the labeled substrate but not be detected and (6) abundant organisms can be found in all density fractions as demonstrated here and previously (Youngblut, Barnett, and Buckley 2018), hence they may be erroneously classified as incorporators when sequencing only heavy fractions.

We propose that sequencing all fractions should overcome all of these obstacles. A study demonstrating the feasibility of this approach using qSIP with 3 density fractions in soil has been published recently (Starr et al. 2018). The use of low gradient resolution limits detection to highly enriched taxa. However, medium gradient resolution along with the decreasing price of shotgun sequencing should still keep the financial and computational costs of MG-SIP manageable while maintaining the detection limit achievable at high resolution. Specifically, our data suggests that circa 10 density fractions (fraction size 0.011 g ml^-1^) the increase in error compared to higher resolution is minor.

All commonly used SIP protocols rely on a linear conversion between mean weighted density of the unlabeled genome and its GC content (Schildkraut, Marmur, and Doty 1962; Buckley et al. 2007; Neufeld et al. 2007; T. Lueders 2010; Murrell and Whiteley 2010). Once again, this inherent variation implies that GC content cannot be accurately converted to density with a canonical equation (Schildkraut, Marmur, and Doty 1962). Rather, we may need to create a calibration curve of %GC/WMD per tube by using an internal standard, as these equations have a very high R^2^ but with varying slopes and intercepts. An accurate conversion between MWD and %GC may become extremely important for SIP experiments in which metagenomes are sequenced only from the heavy fractions of labeled samples. Once genomic bins are assembled, their %GC can be converted into a theoretical unlabeled WMD which can be used to calculate the density shift, and thus the enrichment level of those bins. A reliable calculation may allow us to avoid analyzing most of the unlabeled controls, and thus save on labor and costs To demonstrate the costs of high-resolution MG-SIP, we compared the resources and yield of a simplified experiment. High resolution SIP routinely generates 40-60 density fractions.

Assuming the conservative number of 40 fractions, we would generate 40 metagenomes from each tube. As we would sequence not only the isotopically labeled fractions but also the control fractions, we would be looking at 80 metagenomes. Furthermore, for a minimum of 3 replicates, the number would increase to 240 metagenomes. Assuming shallow sequencing of 2 Gbp per metagenome, the sequencing process would yield 480 Gbp that would need to be stored, manipulated and assembled. Using the latest NovaSeq platform that produces higher yield at a lower cost per-base, we would still require 5 lanes on the sequencer. The cost of these combined with library preparation would currently revolve around $50,000 (http://qb3.berkeley.edu/gsl/wp-content/uploads/2018/08/2018_2019-QB3-Genomics-Rates_August.pdf). In addition, there would be a cost in labor or robot facility time for fractionation, precipitation, ethanol washing and elution. As stated, this would be a highly simplified experiment. All of these costs would increase when adding time-points or other experimental conditions such as different temperatures, other nutrients etc.

Reducing the number of fractions to 10 would yield a relative error lower than 0.0005 g ml^-1^ which is negligible considering that the standard deviation of the WMD is at least 3 times higher than that even when using an automated pipeline. Below a fraction size of 0.011 g ml^-1^ the mean relative error increase, as does its variability, in both replicated and unreplicated datasets.

However, this increase in variability can still be mitigated by reallocating some of the funds towards replication. In fact, reducing the number of fractions even by a factor of 2 will allow for doubling the number of replicates without additional costs, while increasing the statistical power of any downstream analyses.

With MG-SIP we would, in theory, already have the GC content of a genomic bin, so that it would not need to be calculated from the mean weighted density, in which case the control would be used only to calculate the density shift. That being said, we expect that the genomic bins generated from metagenomes will not be complete, therefore they may also have some error rate in GC content calculation. When combining MG-SIP with an internal %GC/density ladder, we could significantly decrease the number of unlabeled controls sequenced and use the calibration curve from the labeled tubes with the GC content of the bin to calculate the unlabeled WMD and the density shift. The internal standard should be easily informatically separable from the sample. This could be done by creating a mock community of organisms which are highly unlikely to appear in the sample (e.g. in a soil sample use genomes of strictly marine organisms) and could be customizable per experiment. Due to the variation within spin, an external ladder (a mock community in a separate ultracentrifuge tube) would be insufficient. However, it could be argued that finding a suitable set of non-indigenous genomes distinguishable from a highly diverse environment such as soil may prove difficult. Alternatively, if highly complete genomic bins can be assembled, then their %GC would be more reliable, and their WMD can be calculated from the gradient. Such bins could be used as an internal standard for generating a WMD-to-GC formula. As the generation of high-completion bins could only be assessed post hoc, we would still recommend the use of an internal standard.

The inherent variability in qSIP can stem from many steps along the way: replicate variation, bottle effects during incubation, extraction efficiencies, tube to tube variation in the gradient, amplification bias, strain heterogeneity, among-treatment shifts in community composition, and OTU clustering errors (for marker genes). Quantifying the sensitivity of qSIP to those factors will improve existing amplicon-based qSIP techniques and facilitate efficient ways of extending SIP to more ambitious applications, such as metagenome-assembled genome-based SIP.

## Acknowledgements

The authors would like to thank the SPRUCE team, Michaela Hayer, Paul Dijkstra, Sheryl Bell and Ember Morrissey for generously providing data, and Andy Tomatsu at the DOE Joint Genome Institute for operating the high throughput SIP pipeline for the mock communities. Support for analyses and data integration was provided by the U.S. Department of Energy, Office of Biological and Environmental Research, Genomic Science Program LLNL ‘Microbes Persist’ Scientific Focus Area (award #SCW1632), and awards DE-SC0016207 and DE-SC0020172 at Northern Arizona University. Work conducted at LLNL was contributed under the auspices of the US Department of Energy under Contract DE-AC52-07NA27344, and at the Lawrence Berkeley National Laboratory through Contract No. DE-AC02-05CH11231.

